# A mathematical model of pathology progression in the TgF344-AD rat model of Alzheimer’s disease

**DOI:** 10.64898/2026.01.23.701333

**Authors:** Micah Hesketh, Peter Hinow

## Abstract

Alzheimer’s disease (AD) is a devastating neurodegenerative disease whose etiology is poorly understood and for which current treatments provide only modest control of symptoms. To better investigate the causes and progression of the disease, the transgenic TgF344-AD rat model has emerged as a crucial tool. In this paper, we collect observations on the accumulation of amyloid-*β*, changes in neuronal density, and a decline in cognitive performance in TgF344-AD and wild-type rats. We develop a compartmental ordinary differential equation model and determine its parameters by fitting the output to the experimental observations in the literature. Our model simulations are compatible with the hypothesis that the accumulation of amyloid-*β* leads to a rapid decline in neuronal density followed by a significant loss in memory and learning ability. Our mathematical model can provide a bridge between AD research in rodent models and the human condition of AD.

## 1 Introduction

Animal models of human diseases have long proved their usefulness in improving our understanding of the original pathologies in our species. Although many challenges remain to translate the knowledge gained in, for example, small rodents to *Homo sapiens* (Ineichen et al., 2024), research continues to characterize pathologies, identify targets, and evaluate new therapeutic agents and treatments (McGonigle and Ruggeri, 2014; Loewa et al., 2023). A particular field of interest is progressive neurodegenerative diseases in humans, such as old-age dementia. Alzheimer’s disease (AD) is a devastating age-related neurodegenerative disease that currently affects approximately 50 million people worldwide. This number is expected to triple by 2050, creating enormous strains for caregivers and financial burdens on healthcare systems. In this paper, we propose a mathematical model to describe the progression of the pathology in the transgenic rat model TgF344-AD (Cohen et al., 2013) of AD and thus combine two different methods of disease modeling. The TgF344-AD rat model is unique in that it represents all hallmarks of human AD such as progressive amyloid-*β* (A*β*) deposition, tauopathy, gliosis, neuronal loss, and cognitive impairment, while expressing only two mutant human genes (Fowler et al., 2022).

The root causes of AD are debated and there are multiple competing and overlapping hypotheses that explain the condition. These hypotheses include the accumulation of metabolic waste and a decrease in neurotransmitters. We will focus our attention on the idea that the accumulation of metabolic waste is the main driver of the progression of AD. Postmortem studies have found that AD patients have significantly higher levels of two types of proteins, A*β* and misfolded tau proteins. Thus, the two natural basic hypotheses of AD are the A*β* cascade and the tau pathology hypothesis. We will focus our attention on the A*β* cascade which proposes that the accumulation of two types of A*β*, A*β*40 and A*β*42 results in insoluble plaques in the brain triggering neuronal degeneration. Both forms of A*β* are produced in healthy brains when amyloid precursor protein (APP) is broken down and cleared by apolipoprotein (ApoE). Agerelated degradation of the glymphatic system and consequent failure of the clearance mechanism for A*β* and other metabolic waste products has been implicated as root cause of AD and other dementias (Nedergaard and Goldman, 2020).

Mathematical modeling of Alzheimer’s disease (and other neurodegenerative diseases) is nowadays an established field; see (Hao and Friedman, 2016; Bertsch et al., 2021; Lindstrom et al., 2021; Hao et al., 2022; Bertsch et al., 2023; Brennan et al., 2024; Thompson et al., 2024; Moravveji et al., 2024; Chu et al., 2024) and the works cited therein. A major challenge remains in linking the variables of the mathematical models with experimental observations, whether they are in humans or model animals. Here, we set out to create a simple ordinary differential equation (ODE) model based on the A*β* cascade hypothesis and draw on a rich set of observations in rats. Transgenic TgF344-AD rats show large accumulations of A*β* in the brain along with measurable decreases in neuronal density in key brain regions followed by behavioral deficits compared to wild-type rats (Cohen et al., 2013; Saré et al., 2020; Bernaud et al., 2022; Fang et al., 2025). Our model aims to connect and quantify these observations. Later, this model can be adapted to incorporate neurofibrillary tangles of misfolded tau protein, possible treatments such as A*β*-targeting drugs or transcranial ultrasound.

This paper is organized as follows. In Section 2 we begin by stating our conceptual model of the A*β* cascade leading to Alzheimer’s disease and follow up with a construction of a three-stage mathematical model formulated as a system of ODEs. In Sections 3 and 4 we collect experimental observations about disease progression in the transgenic rat model TgF344-AD (Cohen et al., 2013) of Alzheimer’s disease. We are particularly interested in A*β*-deposits, changes in neuronal density, and a decrease in learning and memory abilities. We use these published data to parameterize and simulate our mathematical model (no new experimental data were generated in this study). The discussion is contained in Section 5.

## 2 Building the mathematical model

The conceptual basis of our AD model is that the root cause of the condition is the accumulation of harmful proteins; see Figure 1. This type of AD model is in line with, for example, the work in (Hao et al., 2022; Bertsch et al., 2023) and many others. Although we assume the A*β* cascade hypothesis, our model is open to future refinements, such as the inclusion of tauopathy.

**Figure 1:**
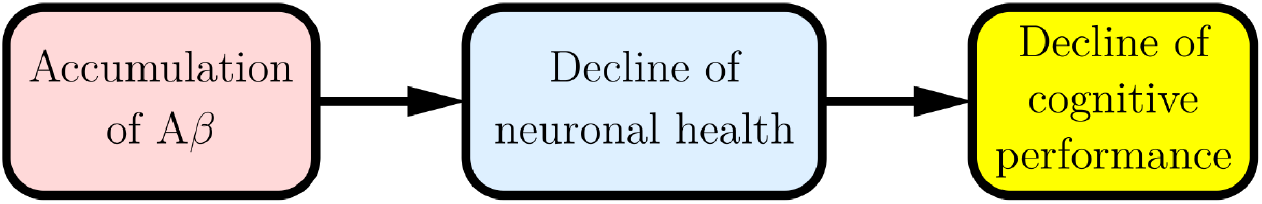
Conceptual model of progression of Alzheimer’s disease.

As the first variable, we denote by *x*(*t*) the amount of neurotoxic A*β* oligomers uniformly distributed in the brain. Following reports by (Jack et al., 2013; Whittington et al., 2018) that A*β* reaches a plateau in humans and the modeling choice of (Hao et al., 2022), we use a logistic ordinary differential equation

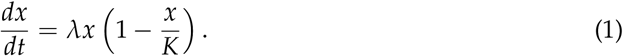

Here *λ* is the rate of formation of A*β* and *K >* 1 is the carrying capacity. The initial condition at *t*_0_ = 6 months is set at *x*(*t*_0_) = 1 representing the normalized amount of A*β* in the brain of a 6 month old TgF344-AD rat. In a Fischer 344 wild-type rat it is set to *x*(*t*_0_) = 0.4 following (Fang et al., 2023, Figure 1 B) .

The equation for neuronal density *y* is

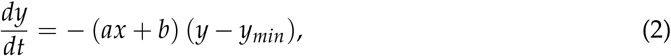

where *a, b* and *y*_*min*_ are positive constants. The constants *a* and *b* describe the decrease in neuronal density that is dependent and independent of A*β* accumulation, respectively. The latter thus can be seen as the natural aging process of wild-type Fischer 344 rats. The constant 0 ≤ *y*_*min*_ *<* 1 represents the minimal neuronal density required for the rat brain to still function. We normalize the experimental neuronal density data with respect to 6-month-old rats such that *y*(*t*_0_) = 1 and *y*(*t*_0_) ≈ 1 for the wild-type and TgF344-AD populations, respectively. This normalization reflects observations from (Cohen et al., 2013) that TgF344-AD rats initially have very similar neuronal density in the hippocampus and cortex as wild-type rats; for more details, see Section 3 and Figure 3 (A) below. The benefit of normalizing the data in this way is that we can monitor the neuronal density of the TgF344-AD rats relative to the “high water mark” and therefore we can observe their decline relative to healthy natural aging.

The impact of losing neurons and synapses may manifest itself only with delay. Defining an abstract mental performance *z*, consisting of memory and cognition abilities, we postulate a third differential equation,

**Table 1:** Sequentially fitted parameters for TgF344-AD and wild-type populations. The 95% confidence intervals for the TgF344-AD population were calculated using the bootstrap method.

| parameter | TgF344-AD | 95% confidence interval | WT |
| --- | --- | --- | --- |
| $\lambda$ ($\text{mo}^{-1}$) | $3.24 \cdot 10^{-1}$ | $[3.12 \cdot 10^{-1}, 3.43 \cdot 10^{-1}]$ | $3.09 \cdot 10^{-1}$ |
| $K$ (au) | 62.34 | [59.03, 66.04] | 5.7 |
| $a$ ($\text{mo}^{-1}$) | $4.2 \cdot 10^{-2}$ | $[3.86 \cdot 10^{-2}, 4.63 \cdot 10^{-2}]$ | $1.78 \cdot 10^{-1}$ |
| $b$ ($\text{mo}^{-1}$) | $7.15 \cdot 10^{-4}$ | $[6.65 \cdot 10^{-4}, 7.49 \cdot 10^{-4}]$ | $1.41 \cdot 10^{-3}$ |
| $y_{min}$ (au) | $5.04 \cdot 10^{-1}$ | $[5.01 \cdot 10^{-1}, 5.06 \cdot 10^{-1}]$ | $8.66 \cdot 10^{-1}$ |
| $c$ ($\text{mo}^{-1}$) | $1.67 \cdot 10^{-1}$ | $[1.30 \cdot 10^{-1}, 2.21 \cdot 10^{-1}]$ | $2.6 \cdot 10^{-1}$ |

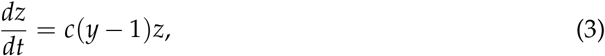

where *c* is a positive constant. As with neuronal density, we normalize behavioral data to 6-month-old rats, so *z*(*t*_0_) = 1 for the wild-type population, and data from (Saré et al., 2020) and (Cohen et al., 2013) indicate *z*(*t*_0_) ≈ 1 for the TgF344-AD population. Note that both *y* and *z* are always decreasing, i.e. declines in neuronal density and mental performance are irreversible.

## 3 Observations in rats and implementation of the minimization procedure

A defining characteristic of TgF344-AD rats is that they have high levels of A*β* at an early age (Fang et al., 2023; Futácsi et al., 2025) compared to wild-type rats. In addition, A*β* accumulates rapidly in TgF344-AD rats (Saré et al., 2020; Bernaud et al., 2022). Let *T*_*x*_ denote the combined set of times for which A*β* data 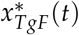 is reported in (Cohen et al., 2013, Figure 3D_2_), (Saré et al., 2020, Figure 6), and (Bernaud et al., 2022, Figure 8B). These measurements are Western blots, A*β* plaque areas or Thioflavin-S burdens from the hippocampus or various areas of the cortex. The amount of A*β* in wild-type rats is much more difficult to discern than for the TgF344-AD animals since many measurements are not carried out for the control group. Nevertheless, it is possible to see a weak A*β* signal in the wild-type population, namely in (Cohen et al., 2013, Figure 4 F-H) that we translate into numerical values in our Figure 2 (A). Furthermore, let *T*_*y*_ denote the set of times for which neuronal density data are reported in (Cohen et al., 2013, Figure 8C), and *T*_*z*_ the set of times for which behavioral data is reported in (Cohen et al., 2013, Figure 2) and (Saré et al., 2020, Figure 1). The mean of these data at time *t* is denoted by (*x*/*y*/*z*)^∗^(*t*) and (*x*/*y*/*z*)(*t*) are the numerical solutions of equations (1)-(3).

**Figure 2:**
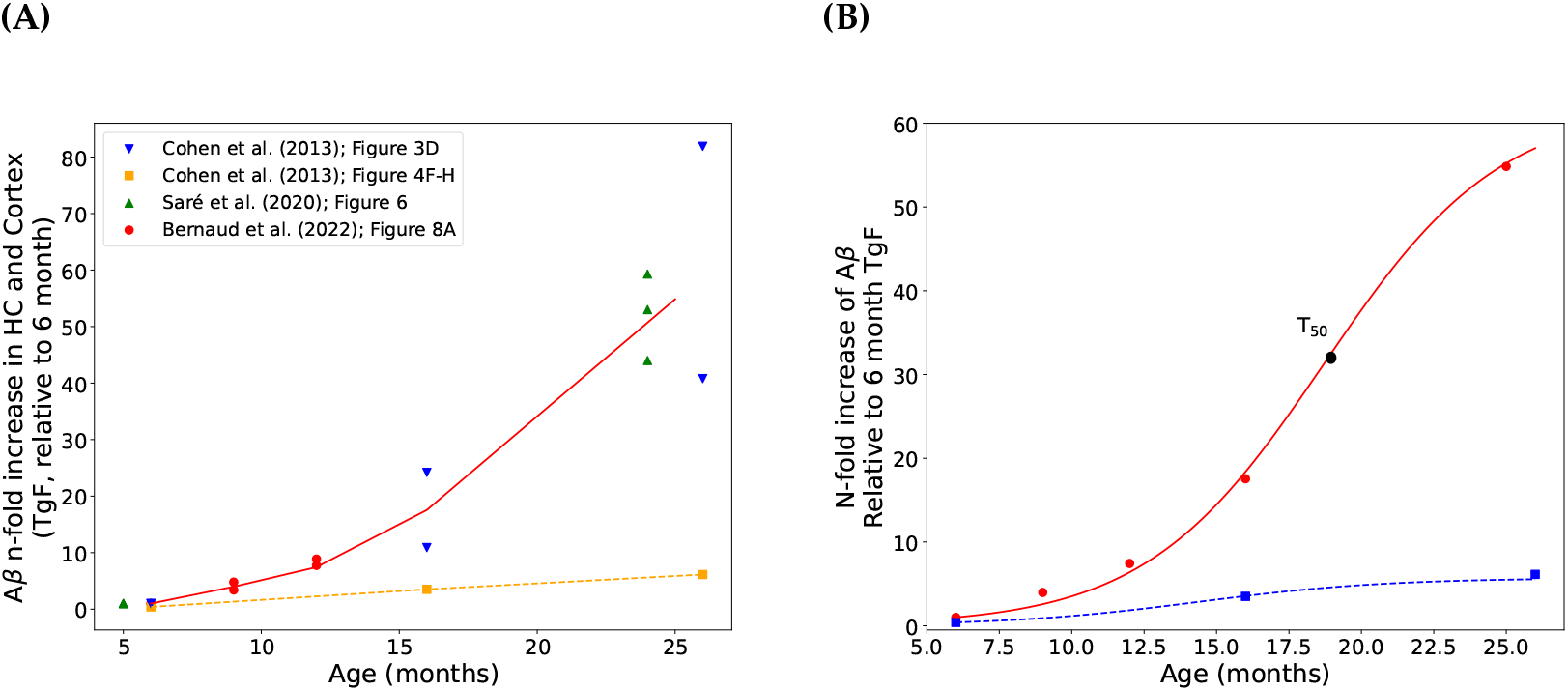
Accumulation of A*β* in the brain of TgF344-AD rats. **(A)** Accumulation of A*β* in the brain of TgF344-AD and wild-type rats. Sources of data are (Cohen et al., 2013, Figure 3D, cortex or hippocampus, blue triangles), (Cohen et al., 2013, Figure 4G, wild-type, orange squares) (Saré et al., 2020, Figure 6, cortex and hippocampus, green triangles), and (Bernaud et al., 2022, Figure 8B, dorsal hippocampus, red dots). The red line represents the average of all TgF344-AD data 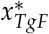. **(B)** The red curve shows the optimal fit of the mean of the data in panel A with equations (1)-(3); shown is the *x*-component. The black dot indicates the inflection point at *T*_50_ = 19 months. The blue dashed line shows the level of A*β* in the brain of wild-type rats.

Due to the sequential nature of the pathology and its mathematical model, we implement a sequential fitting procedure. We determine the parameter vector *θ* = (*λ, K, a, b, c, y*_*min*_) by minimizing first

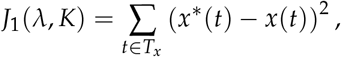

followed by minimizing

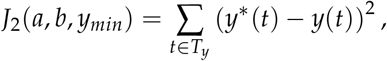

and finally

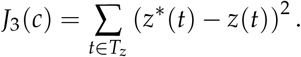

We find that a joint fitting of the parameter vector *θ* results in similar parameter values. These are shown in the Supplementary Information.

While data on A*β* deposition and neuronal density can be easily extracted from the sources cited, the same is not true about the cognitive performance data. Many of the tests have differing units, for example, the time taken to complete a task or the number of errors made. To make data from different sources comparable, we normalize all data from each trial to the wild-type performance at 6 months. In the Barnes maze experiments in (Cohen et al., 2013, Figure 2 H & I), the decreasing number of errors on days 1, 2, 3, and 4 is translated into the negative slope of the learning process that is then compared for the different groups. (Saré et al., 2020, Figure 1) report working memory measured through a reward-alternating T-maze. For our purpose, we pooled the male and female subgroups.

The ODEs (1)-(3) are numerically solved using a fourth-order Runge-Kutta method (Burden et al., 2018). Each minimization is performed in Python utilizing the scipy implementation of the Nelder-Mead algorithm (Press et al., 2007; Virtanen et al., 2020). The generation of code was accelerated by the use of Google’s Gemini 3.1 Pro (Gemini Team, 2023); however, the results were carefully inspected. The data extracted from the earlier publications and the Python codes will be available from figshare.com/articles/software/ /32690889.

## 4 Results

The combined list of optimal parameter values is given in Table 1 with the numerical solutions shown in panels (B) of Figures 2-4. It is important to note that these values depend on our assumption that the amyloid-*β* cascade is the sole driver of neurodegeneration and behavioral deficits. Our parameters *λ*_*TgF*_ and *K* imply that the inflection point is located at *T*_50_ = 19 months. Note that the A*β*-independent rate of decline in neuronal density *b* the TgF344-AD population may be an effect of neurofibrillary tangles of misfolded tau protein, another hallmark of the TgF344-AD rat (Cohen et al., 2013, Figure 5).

We have applied the bootstrapping method (Efron and Tibshirani, 1994; Givens and Hoeting, 2013) but only for the TgF344-AD population, since we have high quality A*β* data only for these. To a numerical solution of our model with the standard parameters, we add independent random errors that follow Gaussian normal distributions 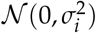. The standard deviations *σ*_*i*_ of the residuals (*i* ∈ {*x, y, z}*) are estimated from the difference between the observed data and the predictions of the best-fit model. Then we fit the model to the artificial data sets and repeat this 2000 times. This allows to determine the 95% confidence intervals, see Table 1. When all parameters are normalized to 1, the widths of the intervals indicate the sensitivity of the model to the corresponding parameter, see Figure 5.

To judge our model against a simpler version, we have set *a* = 0 for the TgF344-AD population and calculated the Akaike Information Criterion (Akaike, 1974) with and without this constraint. For the least square fit we have (up to a constant)

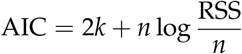

where RSS is the residual sum of squares (i.e. how well the model fits the data), *n* is the number of observations, and *k* is the number of parameters. The result is

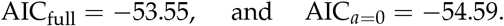

Although this suggests that the restricted model is superior, the small difference ΔAIC *<* 2 makes it difficult to separate them with certainty (Burnham and Anderson, 2002, p. 70). On closer inspection of the data, the neuronal density *y* declines immediately after 6 months whereas *x* needs some time to grow appreciably over its initial value 1. Interestingly, the TgF344-AD transgenic rats show increased levels of anxiety before showing any cognitive decline (Pentkowski et al., 2018; Lopez et al., 2024).

Following the A*β* cascade hypothesis, a natural treatment approach is the removal of builtup A*β*. In (Fang et al., 2025) the authors administer a soluble epoxide hydrolase inhibitor (1-(1-Propanoylpiperidin-4-yl)-3-[4-(trifluoromethoxy)phenyl]urea; TPPU) through the drinking water to six-month-old TgF344-AD rats. They observe

1. an attenuation of A*β* accumulation (Fang et al., 2025, Figure 2 B & C), where A*β*-levels in TPPU-treated TgF344-AD rats at 9-10 months of age are ≈16 % of those in the untreated TgF344-AD rats,
2. neuronal numbers that are equal to the wild-type group at 9-10 months of age (Fang et al., 2025, Figure 8 B & C), and
3. a reduced cognitive impairment compared to the untreated TgF344-AD group at 9-10 months of age (Fang et al., 2025, Figure 8 D).

Our simulation results are shown in Figure 6, where we use *λ*_*TgF*;*TPPU*_ = 1.49 ·10^−1^ mo^−1^ and *K*_*TgF*;*TPPU*_ = 69.99, which are chosen to match the reduction of A*β*-levels at 9-10 months.

## 5 Discussion

In this paper, we have compiled existing experimental data from wild-type and TgF344-AD rats and analyzed them with the help of our three-step model (1)-(3). We find that the A*β* cascade hypothesis is a plausible explanation for the decline in neuronal density and the loss of cognitive performance in TgF344-AD rats. Given the moderate complexity of our model, it provides a satisfactory fit to the experimental data. In addition, data taken from different sources (Cohen et al., 2013; Saré et al., 2020; Bernaud et al., 2022) align to form a coherent picture. We did not select behavioral data from (Bernaud et al., 2022) where the TgF344-AD rats performed worse compared to wild-type rats already at six months of age, and then the discrepancy remained the same at nine and twelve months. This performance gap may be caused by the increased levels of anxiety that transgenic rats are known to have at an early age (Pentkowski et al., 2018; Lopez et al., 2024). The tests conducted by (Bernaud et al., 2022) required the rats to swim to platforms. As rats are not inherently aquatic animals, it stands to reason that this environment could increase their anxiety and thereby reduce the performance of the AD group. The data we chose come from tests conducted outside of water, and we believe reduce anxiety as a potential confounding variable on the rats’ performance.

In general, the field remains in flow and conflicting evidence cannot be overlooked entirely. (Futácsi et al., 2025) show a negative correlation between the number of A*β* plaques in the hippocampus and prefrontal cortex, on the one hand, and the number of various types of neurons, on the other. However, they state that they could not detect any correlation between memory impairment and neuropathological changes in the brain. In another recent work, (Bac et al., 2023) found no significant correlation between amyloid plaques or levels of insoluble A*β* and the learning index even in some aged TgF344-AD rats. Thus, they are led to the conclusion that *“Aβ is not the main contributor to cognitive impairment”*. We consider the original longitudinal data from (Cohen et al., 2013) succinct to build a mathematical model around them. A future extension of the mathematical model may also incorporate the formation of neurofibrillary tangles due to the accumulation of tau protein and their detrimental effect on neuronal density. Apart from (Cohen et al., 2013, Figure 4 H-F), the amount of A*β* in the wild-type control group is reported in (Fang et al., 2023, 2025; Futácsi et al., 2025) but only at a single time point. It should be stated that the value of *a* reported for the wild-type group in Table 1 is probably an overestimation. Rat A*β* differs from human A*β* in three amino acids (Ueno et al., 2014) and does not form the characteristic lesions of human AD. It is therefore unlikely to have the same neurotoxic effect as human A*β*.

In experiments, a slower learning rate can be clearly observed in the Barnes maze (Barnes, 1979) in wild-type rats at 24 months (Cohen et al., 2013, Figure 2I_2_). At the same age, they show only a minimal decline in neuronal density in the hippocampus (Cohen et al., 2013, Figure 8C_2_) and a small decline in neuronal density in the cortex (Cohen et al., 2013, Figure 8C_1_). Some behavioral aspects decline naturally towards the end of the life of the inbred Fisher 344 rat strain. This strain is commonly used in aging studies and forms the basis for the transgenic TgF344-AD model. We are using neuronal density as the only measure of neuronal health. An alternative or complementary choice would be the level of N-acetylaspartate (NAA) in different regions of the brain, whose decline has been associated with neuronal loss and dysfunction (Muñoz-Moreno et al., 2022). There are also various magnetic resonance imaging (MRI) connectome data from TgF344-AD rats (Anckaerts et al., 2019; van den Berg et al., 2022). Significant difficulty remains in their interpretation as indicators of structural and functional brain integrity.

How can our insights be related to research on Alzheimer’s disease in humans? For example, Jack et al. (2013) report that the growth of A*β* reaches a plateau by analyzing positron emission tomography (PET) scans of patients with AD and patients with mild cognitive impairment (MCI). Fitting a logistic function similar to the solution of equation (1) to the PET data (Whittington et al., 2018, p. 825) yields an approximate growth rate in humans of *r* = 0.2 y^−1^. The carrying capacity is approximately *K* = 1.5 with respect to the standard uptake value ratio (SUVr) of the PET agent ^18^F-AV-45 (florbetapir). Furthermore, (Whittington et al., 2018, Figure 3) showed that (PET-) carrying capacities vary between different regions of the brain but are generally between 1 and 1.5. (Whittington et al., 2018) also report a global location of the inflection point *T*_50_ = 15 years, i.e. halfway through the 30-year disease cascade. In Figure 7 we superpose the growth curve from (Jack et al., 2013, Figure 2B) and an affine image of our optimal solution *x*_*TgF*_(*t*) from Figure 2 and Table 1, extended to 26 months after the beginning at 6 months. The affine image is given by

**Figure 3:**
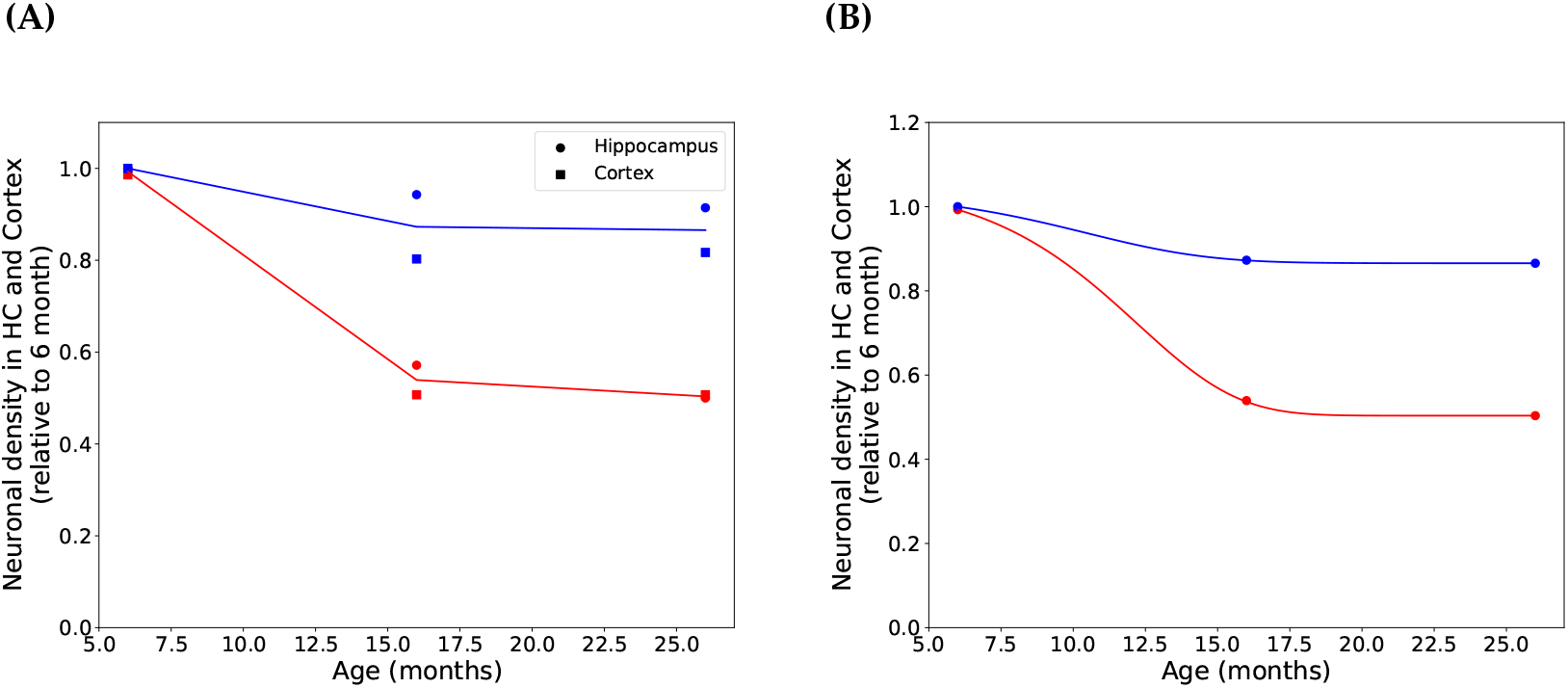
Decline in neuronal density. **(A)** Decline in neuronal density in the hippocampus and cortex of wild-type (blue symbols) and TgF344-AD (red symbols) rats, from (Cohen et al., 2013, Figure 8C). The solid lines indicate the averages of the two regions 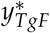 (red) and 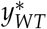 (blue), respectively. **(B)** The *y*-component of the numerical solution to equations (1)-(3) with the parameters from Table 1 is plotted for both populations together with the averaged data from panel A. Blue and red curves and symbols represent wild-type and TgF344-AD populations, respectively.

**Figure 4:**
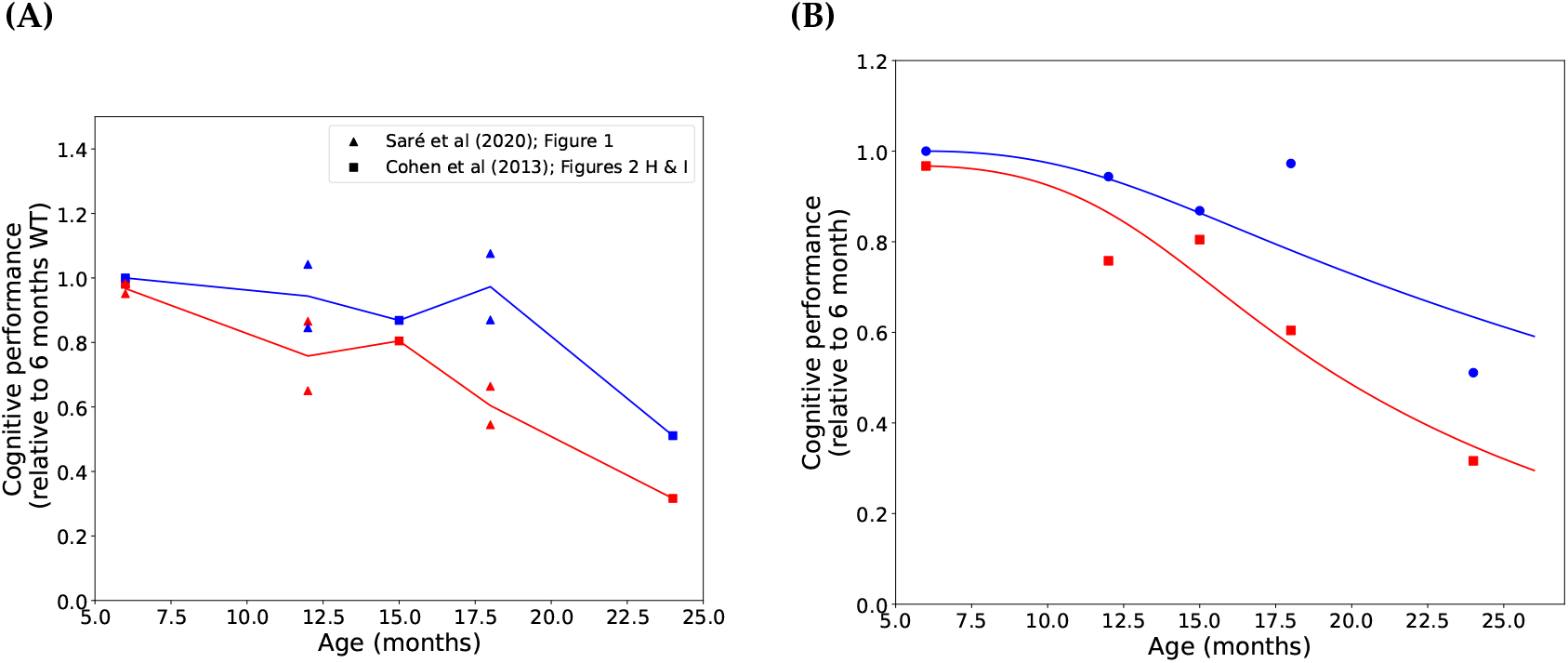
Decline in cognitive performance. **(A)** Relative cognitive performance data reproduced from (Cohen et al., 2013; Saré et al., 2020) normalized to 6 month old wild-type performance. Blue symbols represent wild-type animals and red symbols represent TgF344-AD animals, respectively. The lines represent the mean of the data 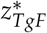 and 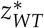 corresponding in color. **(B)** The *z*-component of the numerical solution of equations (1)-(3) fitted to the mean of the data in panel A. Blue and red curves and symbols represent wild-type and TgF344-AD populations, respectively.

**Figure 5:**
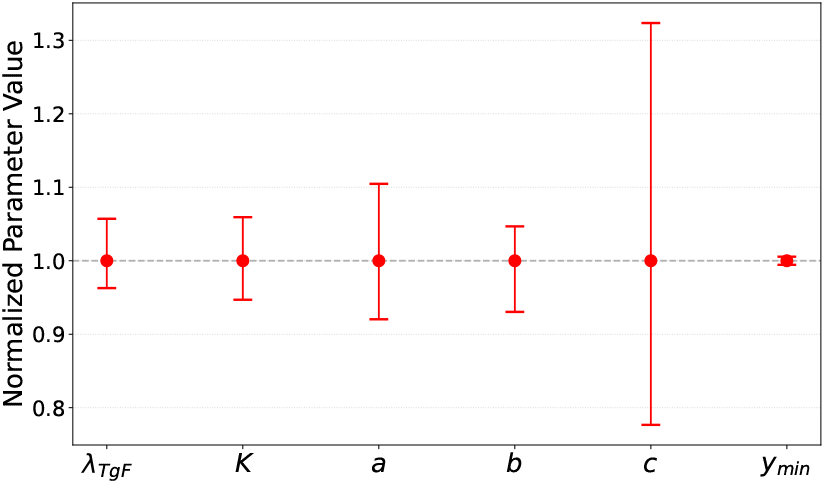
Confidence intervals for parameters for the TgF344-AD population. For comparison, we have normalized all parameter values to 1.

**Figure 6:**
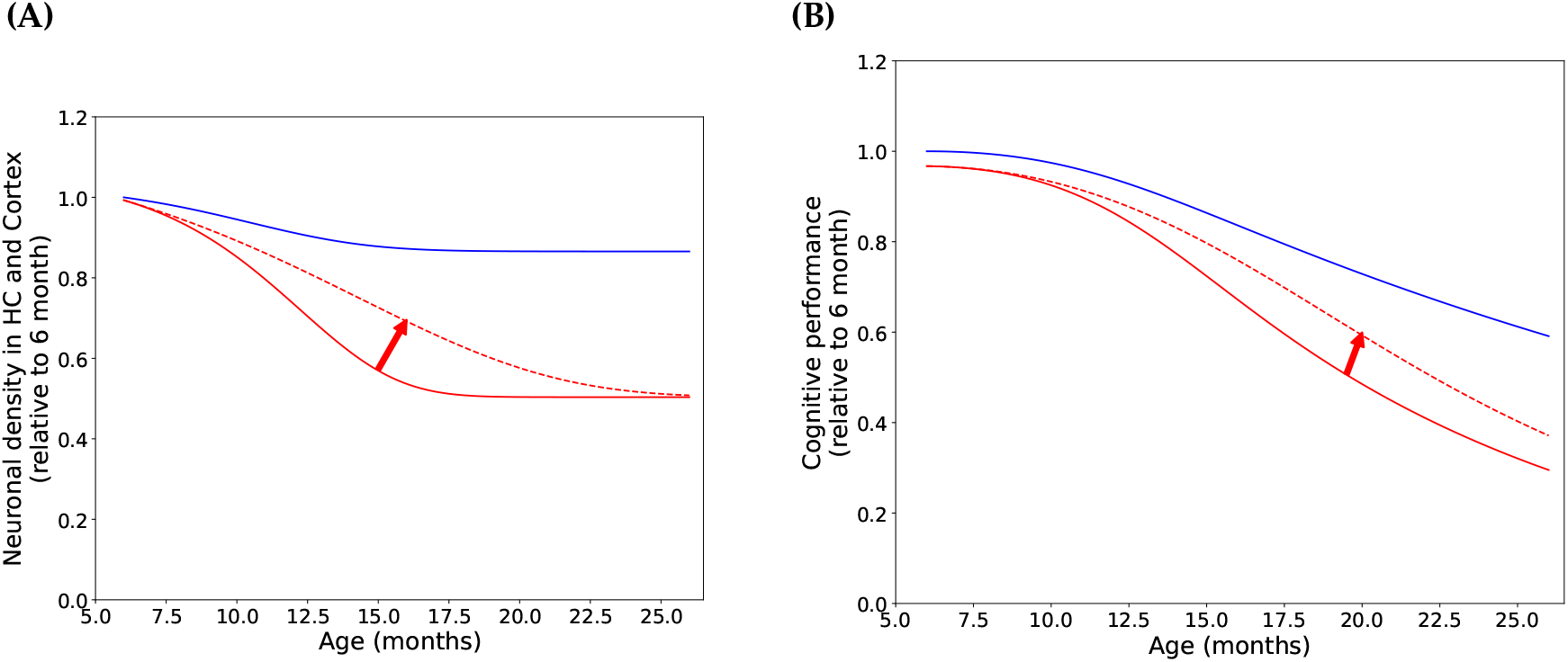
Treatment of TgF344-AD rats with TPPU. **(A)** Simulated slowed down of neuronal density under TPPU treatment (dashed red line). **(B)** Simulated slowed down of cognitive decline under TPPU treatment (dashed red line). The solid curves are taken from Figure 3 and 4, respectively. The red arrows emphasize the improvement.

**Figure 7:**
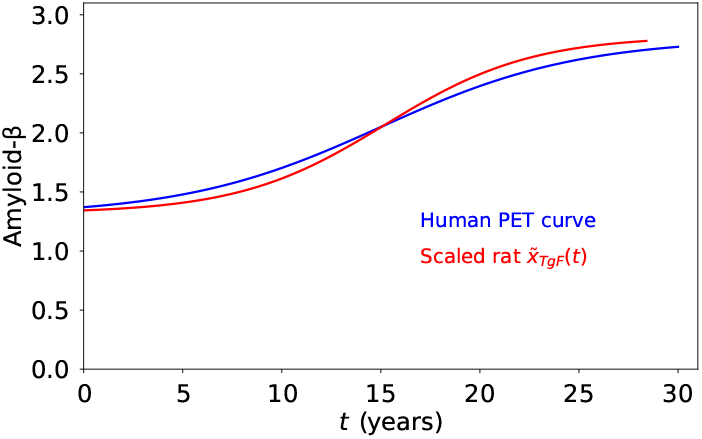
Comparison of human and rat model A*β* cascades. The consolidated human A*β* growth curve (Jack et al., 2013, Figure 2B) and (Whittington et al., 2018, Figure 3) from the PET data (blue) and an affine image of the A*β* accumulation in TgF344-AD rats (equation (4), red). Both curves exhibit approximately the same slope at the inflection point.

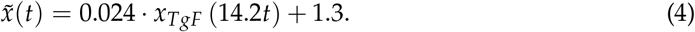

Under this scaling, we find remarkable agreement between the two growth processes in humans and rodents, in particular, the slopes of the two curves at their inflection points.

The fact that A*β* oligomers reduce neuronal functionality is incorporated in numerous mathematical models of human AD analogously to the bilinear term −*axy* in our equation (3). For example, in (Lindstrom et al., 2021, Equation (7)), the spatially dependent “cell viability” *V* with values in [0, 1] is modeled as

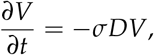

where *D* is the local concentration of A*β* dimers (which is another variable of the model). The damage rate of A*β* dimers is given by *σ* = 4.94 M^−1^ s^−1^ (Lindstrom et al., 2021, Table 1), derived from cell viability assays in mice (Lambert et al., 1998) and rat (Cizas et al., 2010) brain cells. The precise impact of A*β* on human and rodent neuronal tissue will remain a central topic for future research. It is remarkable that A*β*_42_ has been observed at concentrations varying over 12 orders of magnitude (Raskatov, 2019). Petrella et al. (2019) were the first to incorporate the link between neuronal dysfunction and subsequent decline in cognitive and learning abilities, as we do here in our model equation (3). By their very nature, the behavioral symptoms of neurodegenerative diseases are difficult to quantify, making it difficult to compare them to the output of a mathematical model. This is in sharp contrast to, for example, mathematical models of cancer where the number of offending cells readily lends itself as the fundamental variable. Clearly, a link between neuronal density and cognitive ability must exist, but it is much less clear how it should be implemented mathematically. Although the brain is a very sensitive organ, it nevertheless possesses a certain degree of adaptability and plasticity. Mild to moderate neuronal loss in some circuits may be compensated for to at least some extent by other parts. In the case of TgF344-AD rats, regular training and engagement with the animals may reduce anxiety (Lopez et al., 2024) and the impact of declining neuronal health on the cognitive performance (Casanova-Pagola et al., 2026).

## Acknowledgments

The authors thank the Department of Mathematical Sciences at the University of Wisconsin-Milwaukee for support of Micah Hesketh through a graduate fellowship. We are indebted to Dr. Muna Aryal (North Carolina Agricultural and Technical State University, Greensboro, NC) for discussions that formed the starting point of this work. We thank two unknown readers for valuable comments that helped to improve the manuscript.

## Competing interests

The authors declare no competing interests.

## Notes

### Competing Interest Statement

The authors have declared no competing interest.

### Summary of Updates

The manuscript has been revised following the recommendations of two reviewers.

