## Supplemental table and figure for "A mathematical model of pathology progression in the TgF344-AD rat model of Alzheimer’s disease"

### Supplementary Information

Micah Hesketh Peter Hinow

August 26, 2026

We have also implemented a joint parameter fitting strategy for the vector $\theta = (\lambda, K, a, b, c, y_{min})$ where we use a cost functional

$$J(\theta) = \sum_{t\in T_x} \left( \frac{x^*(t) - x(t)}{\widehat{x^*}} \right)^2 + \sum_{t\in T_y} (y^*(t) - y(t))^2 + \sum_{t\in T_z} (z^*(t) - z(t))^2 .$$

Here

$$\widehat{x^*} = \max_{t\in T_x} x^*(t)$$

is used to normalize the A$\beta$ data to the same scale as the rest. The results are presented in Table 1. The differences between the two strategies are small for most parameters in the two populations. The differences in the values of the fitted parameters in the wild-type population can be ascribed to the fact that the A$\beta$ data were not obtained from the same experiments as in the TgF344-AD group. It is noticeable, however, that the parameter $b$, the A$\beta$-independent rate of neuronal density decline is almost two orders of magnitude larger for the TgF344-AD population under the joint fitting strategy. This has an implication on the predicted effect of a reduction of $\lambda$ through therapy, within the ramifications of the present model.

| parameter | TgF344-AD | WT |
|---|---|---|
| $\lambda$ (mo$^{-1}$) | $3.24 \cdot 10^{-1}$ | $3.09 \cdot 10^{-1}$ |
| $K$ (au) | 62.34 | 5.7 |
| $a$ (mo$^{-1}$) | $2.21 \cdot 10^{-2}$ | $7.36 \cdot 10^{-3}$ |
| $b$ (mo$^{-1}$) | $5.58 \cdot 10^{-2}$ | $2.44 \cdot 10^{-2}$ |
| $y_{min}$ (au) | $4.87 \cdot 10^{-1}$ | $7.67 \cdot 10^{-1}$ |
| $c$ (mo$^{-1}$) | $1.64 \cdot 10^{-1}$ | $3.71 \cdot 10^{-1}$ |

Table 1: Jointly fitted parameters for TgF344-AD and wild-type populations.

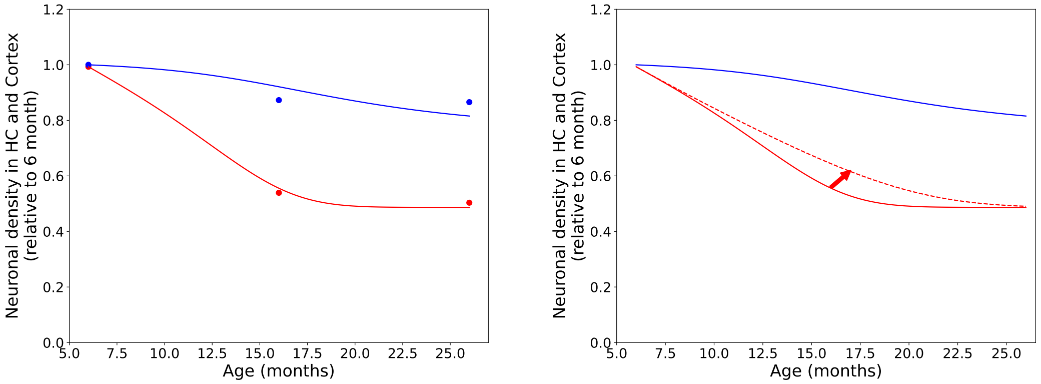


Figure 1: **Decline in neuronal density.** **(Left)** Evolution of neuronal density (blue for wild type, red for TgF344-AD) using the joint fitting strategy. **(Right)** Same as left, but with the TPPU simulation (dashed line).

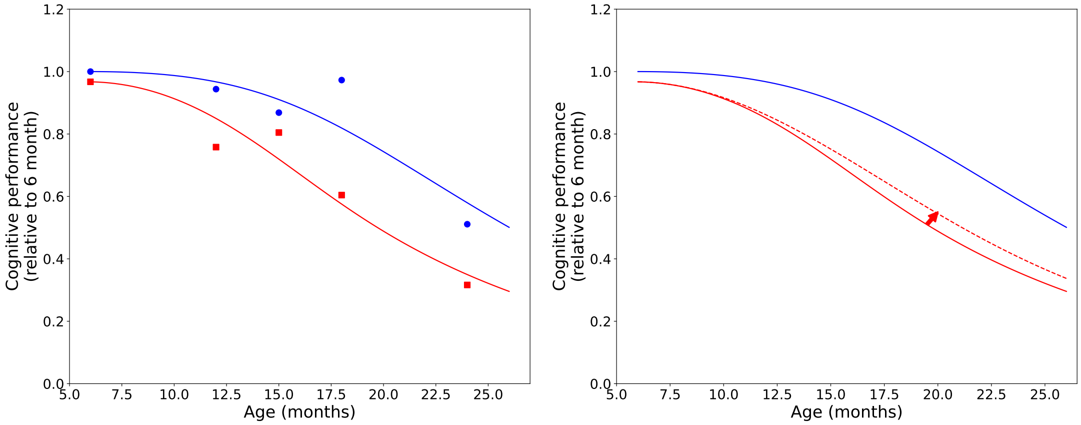


Figure 2: **Decline in neuronal density.** **(left)** Evolution of cognitive performance (blue for wild type, red for TgF344-AD) using the joint fitting strategy. **(Right)** Same as left, but with the TPPU simulation (dashed line).
